# Stabilized dengue virus 2 envelope subunit vaccine redirects the neutralizing antibody response to all E-domains

**DOI:** 10.1101/2024.07.18.604114

**Authors:** Devina J. Thiono, Demetrios Samaras, Thanh T.N. Phan, Deanna R. Zhu, Ruby P. Shah, Izabella Castillo, Lawrence J. Forsberg, Lakshmanane Premkumar, Ralph S. Baric, Shaomin Tian, Brian Kuhlman, Aravinda M. de Silva

## Abstract

The four-dengue virus (DENV) serotypes cause several hundred million infections annually. Several live-attenuated tetravalent dengue vaccines (LAVs) are at different stages of clinical testing and regulatory approval. A major hurdle faced by the two leading LAVs is uneven replication of vaccine serotypes stimulating a dominant response to one serotype at the expense of the other three, leading to the potential for vaccine antibody (Ab) enhanced more severe infections by wild type DENV serotypes that fail to replicate in the vaccine. Protein subunit vaccines are a promising alternative since antigen dosing can be precisely controlled. However, DENV envelope (E) protein subunit vaccines have not performed well to date, possibly due to differences between the monomeric structure of soluble E and the E homodimer of the viral surface. Previously, we have combined structure-guided computational and experimental approaches to design and produce DENV2 E antigens that are stable homodimers at 37℃ and stimulate higher levels of neutralizing Abs (NAbs) than the WT E antigen in mice. The goal of this study was to evaluate if DENV2 E homodimers stimulate NAbs that target different epitopes on E protein compared to the WT E monomer. Using DENV4/2 chimeric viruses and Ab depletion methods, we mapped the WT E-elicited NAbs to simple epitopes on domain III of E. In contrast, the stable E homodimer stimulated a more complex response towards all three surface-exposed domains of the E protein. Our findings highlight the impact of DENV2 E oligomeric state on the quality and specificity of DENV NAbs, and the promise of DENV E homodimers as subunit vaccines.

**Importance:** The ideal dengue virus (DENV) vaccine should elicit a balanced and highly protective immune response against all 4 DENV serotypes. Current tetravalent live-attenuated DENV vaccines have faced challenges due to uneven replication of vaccine virus strains stimulating a strong immune response to one serotype and weak responses to the other three. Protein subunit vaccines provide novel opportunities to stimulate a balanced response because dosing can be precisely controlled and independent of vaccine virus replication. Here, we compare immune responses elicited by a new DENV serotype 2 protein vaccine designed to match the structure of proteins on the viral surface. We find that proteins designed to match the viral surface stimulate better immune responses targeting multiple sites on the viral surface compared to previous protein vaccines. Our results justify further testing and development of these second-generation DENV protein subunit vaccines.

## Introduction

The four dengue virus (DENV) serotypes are mosquito-borne flaviviruses that are estimated to infect several hundred million people per year living in tropical and sub-tropical regions of the world(1). From January to June 2024, there has been around 10 million suspected dengue cases in the Americas alone(2). Clinically, DENV infections can be asymptomatic or present as dengue fever, dengue hemorrhagic fever and shock syndrome. Vaccination is one of the most effective methods for preventing flavivirus infections and effective vaccines have been developed against other deadly flaviviruses, such as yellow fever virus and Japanese encephalitis virus. DENV vaccines must be tetravalent because immunity to a single serotype has been linked to antibody-enhanced replication of heterologous serotypes, associated with more severe clinical disease(3). Protein subunit vaccination is a well-established technology for developing safe and effective monovalent and multivalent vaccines. However, DENV subunit vaccines based on the viral envelope (E) protein have performed poorly in pre-clinical and early clinical studies(4–8). One plausible explanation lies in the structure of soluble E protein under physiological conditions. While soluble E protein is in equilibrium between monomer and dimer states, at 37°C, the equilibrium favors the monomer that does not display key dimer dependent antibody epitopes on the viral surface recognized by human neutralizing antibodies (NAbs)(8, 9). Using structure-guided computational methods and experimentation, we have produced stable DENV E homodimers that display quaternary structure epitopes recognized by human NAbs(10, 11). Here, we sought to map the specificity of NAbs elicited by DENV2 WT and dimer-stabilized recombinant E protein (rE) in mice.

The most advanced DENV vaccines are based on tetravalent live attenuated virus formulations. While the two leading candidates (Dengvaxia and Qdenga) have been approved for use by some countries in restricted populations, both candidates suffer from uneven replication of attenuated vaccine viruses that frequently elicit an effective immunity against one or two serotypes only(12–15). In clinical trials, children with no pre-existing immunity to DENVs who received Dengvaxia were more likely to be hospitalized than unvaccinated children, and this vaccine is currently only approved for use in DENV-immune children(16). In seronegative children, Qdenga was efficacious against DENV1 and 2 but not DENV3(17). The number of DENV4 cases was insufficient to assess efficacy against this serotype. Given the challenges faced in developing tetravalent live DENV vaccines, we are focusing on subunit vaccines, which are safer and easier to formulate as a balanced tetravalent vaccine since immunogenicity is independent of viral replication.

The E protein is a membrane-anchored structural protein covering the surface of flaviviruses and the main target of neutralizing and protective antibodies (Ab). On the viral surface, 90 E homodimers are tightly packed to create a smooth protein coat with pseudo-icosahedral symmetry(18, 19). Many human Abs that neutralize DENVs and other flaviviruses recognize quaternary structure epitopes that span E homodimers or higher-order assemblies on the virion(20–23). However, soluble DENV rE at physiological temperature (37℃) is mostly monomeric and does not display the native homodimer form(9). The abundance of monomers is potentially a major contributor to the poor performance of DENV subunit vaccines to date.

We previously identified specific amino acid substitutions that stabilized the rE homodimer at 37℃, while preserving the overall structure of the native homodimer, including quaternary epitopes recognized by human NAb(10). We also showed that mice immunized with stabilized DENV2 rE dimer had higher levels of DENV2 NAb than animals immunized with WT rE. In this study, we determined the specificity and epitope targets of the NAbs elicited by the DENV2 rE dimer in comparison to the traditional WT rE when they are used as vaccine antigens in mice.

## Results

### Immunogenicity of stabilized DENV2 recombinant E homodimers: 2 dose study

We performed experiments to characterize the binding and functional properties of Abs stimulated by WT and stable homodimer E protein subunit vaccines. All the recombinant DENV2 E proteins used in the current study have been previously described(10). The protein nomenclature used in the current study and the location of specific mutations are summarized in **Table S1**. Groups of Balb/c mice were primed and boosted (week 3) with WT rE and a stabilized E homodimer (SD*FL) in the presence or absence of an adjuvant (Alum) (**Fig. 1A, B**). The SD*FL antigen has 2 mutations (A259W, T262R) in the E-domain(ED) II at the dimer interface (I2), 2 mutations (F279W, T280P) in the hinge between EDI and EDIII (U6), and a point mutation (G106D) in the fusion loop (I8) that independently contribute to dimer stabilization (10)(**Fig. 1A, Table S1**). At 12 weeks after vaccination, mice immunized with SD*FL had more DENV2 binding Abs compared to animals immunized with WT rE (**Fig. 1C**). We compared the binding of vaccine-induced antibodies to DENV2 rE monomer (M2P4) and SD*FL. As previously described(10), the M2P4 protein has a mutation (G258E) at the dimer interface (M2) that destabilizes the dimer and 3 mutations (S29K, T33V, A35M) in EDI (P4) that increase stability and secretion of the monomer (**Table S1**). SD*FL immunized animals had higher levels of Abs to both E monomer and dimer antigens compared to WT E immunized animals (**Fig. 1D, E**). To compare relative levels of monomer and dimer binding Abs in each group, we calculated the ratio of dimer/monomer binding. SD*FL immunized animals had a response that was skewed in favor of the dimer over monomer compared to WT E immunized animals (**Fig. 1F**). The SD*FL rE antigen also stimulated higher levels of NAbs compared to WT rE (**Fig. 1G**). Collectively, these results indicate that the stable dimer antigen stimulates higher levels of binding Abs (skewed in favor of the E homodimer over the monomer) and NAbs compared to the WT E antigen.

**Figure 1.**
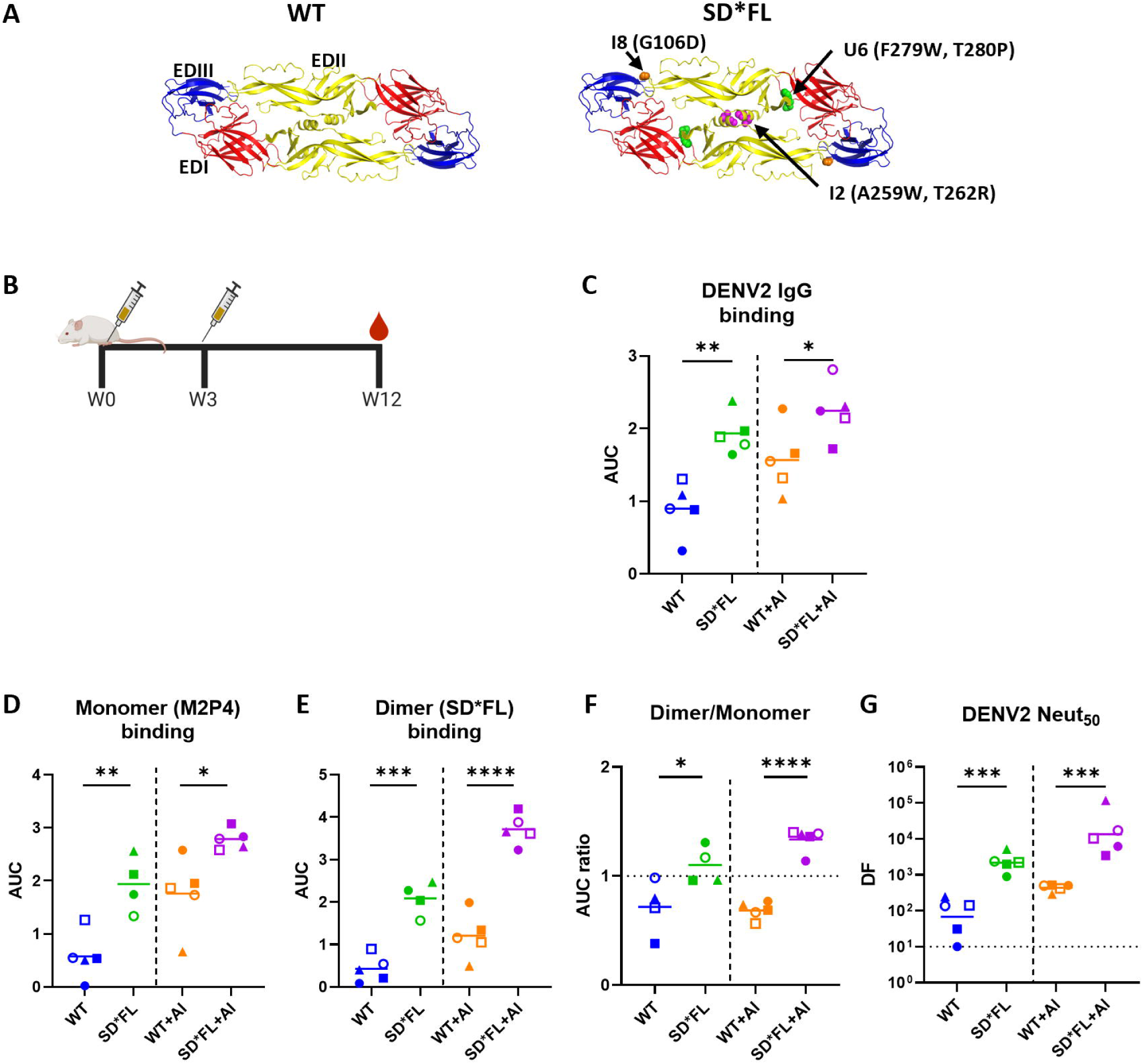
Immunogenicity of recombinant DENV2 E protein vaccines: 2 dose study. (**A**) DENV2 soluble E structure (PDB 1OAN) of the vaccine antigens: wild-type (WT) and SD*FL with stabilizing amino acid mutations shown. (**B**) 6-week old female Balb/c mice were immunized with 5μg of WT or SD*FL rE ±Alum (125ug) at week 0 (W0), and boosted with the same antigen ±Alum at week 3 (W3). IgG binding titer at W12 against (**C**) DENV2, (**D**) DENV2 rE monomer (M2P4), and (**E**) DENV2 rE stabilized dimer with mutated FL (SD*FL) were measured by serially diluting the mouse sera, and the resulting area under the curve (AUC) were reported. Due to limited samples availability, the number of mice in SD*FL group in experiment (**D**) and (**E**) was reduced to n=4. (**F**) To determine the ratio of dimer:monomer binding Ab, AUC value from dimer- and monomer-binding was used. The horizontal line represents the mean titer of all mice within the group. Statistical analysis was done using unpaired t-test (two-tailed, p<0.05). (**G**) DENV2 50% neutralizing Ab titers at week 12. Limit of detection (LoD) for this assay was 1:10. The horizontal line represents the geometric mean titer (GMT) of all mice within the group. Statistical analysis was done on log-transformed values, followed by unpaired t-test (two-tailed, p<0.05). The color and symbols match individual mice tested in different assays.

### Immunogenicity of stabilized DENV2 recombinant E homodimers: 2 vs. 3-dose study

Next, we compared the immunogenicity of 2-dose versus 3-dose vaccination schedules (**Fig. 2A**) in the absence of any adjuvant. Our goal was to obtain moderate to high titer sera required for mapping NAbs stimulated by WT and SD*FL rE antigens. To understand the impact of antigen structure alone on immunogenicity, we did not include the Alum adjuvant, which could alter protein structure(24).

**Figure 2.**
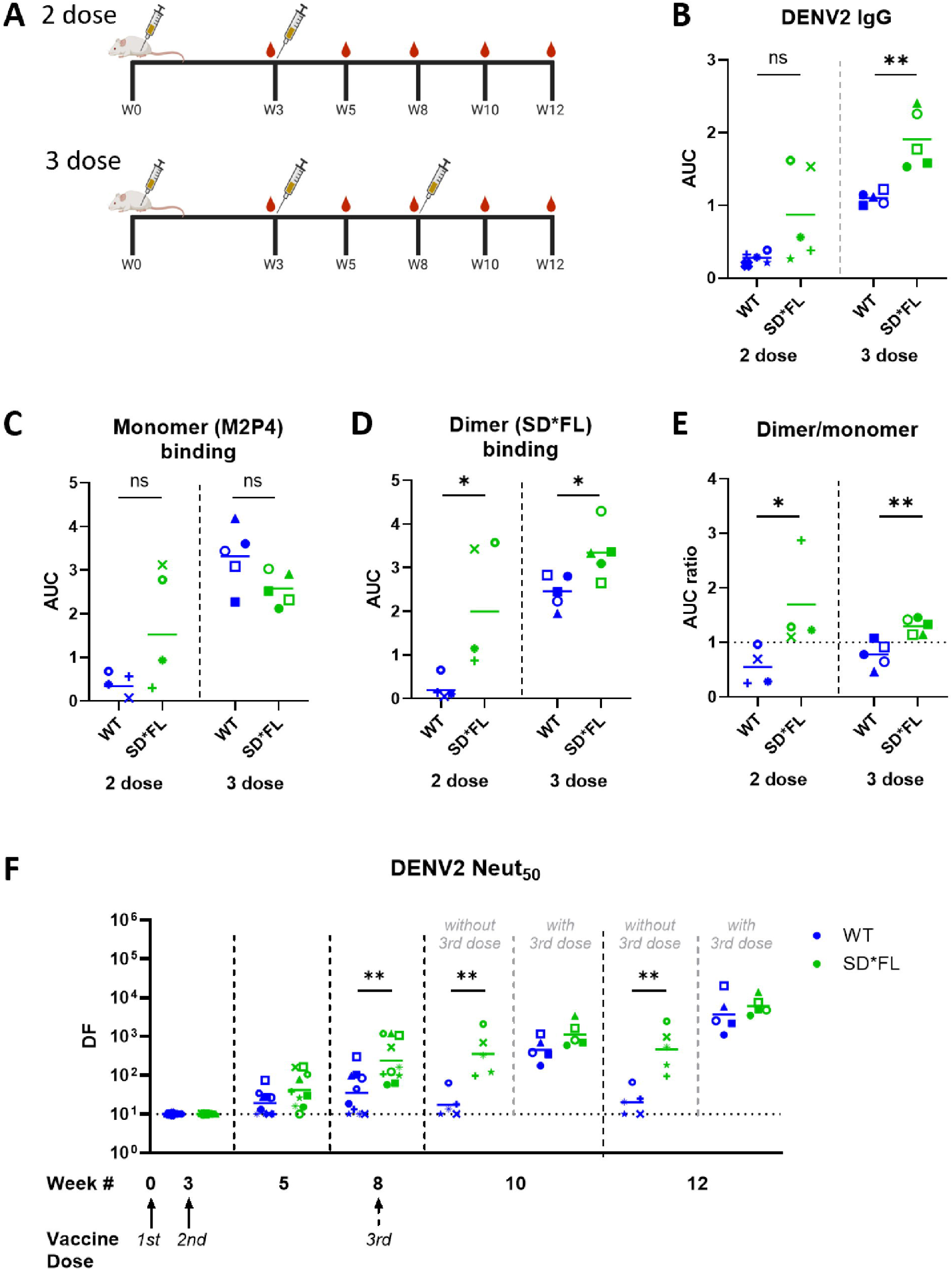
Immunogenicity of recombinant DENV2 E protein vaccines: 3 dose study. (**A**) 6-week old female Balb/c mice were immunized with 5μg of WT or SD*FL rE following two different vaccination schedules: 2 dose schedule-W0 and W3; 3 dose schedule-W0, W3 and W8. Serum samples from each animal were collected pre-(W3, W8) and post-boost (W5, W10, W12). IgG binding against (**B**) DENV2, (**C**) DENV2 rE monomer (M2P4), and (**D**) DENV2 rE dimer with mutated fusion loop (SD*FL) were measured on W12 sera and the resulting AUC was determined. (**E**) We took the AUC value of the monomer and dimer binding to quantify the dimer:monomer Ab ratio. The horizontal line represents the mean titer of all mice within the group. Statistical analysis was done using unpaired t-test (two-tailed, p<0.05). (**F**) DENV2 50% neutralizing Ab titers at week 12. LoD for this assay was 1:10. The horizontal line represents the GMT of all mice within the group. Statistical analysis was done on log-transformed values, followed by unpaired t-test (two-tailed, p<0.05). The color and symbols match individual mice tested in different assays.

The 2-dose component of this study confirmed our previous result (**Fig. 1**) that SD*FL stimulated binding Abs skewed in favor of the dimer and higher levels of NAbs compared to WT rE (**Fig. 2B-E**). As expected, animals in 3-dose groups had higher levels of binding and NAbs than the matched 2-dose groups. In the 3-dose regiment, the SD*FL rE stimulated binding Abs skewed in favor of the dimer over the monomer (**Fig. 2E**). The NAb responses at week 12 were similar for WT and SD*FL rE in the 3-dose study (**Fig. 2F**). Our results demonstrate that in the 2-dose regiment, SD*FL stimulates higher levels of NAbs (compared to WT rE) that are maintained through week 12 (**Fig. 2F**). A third vaccine dose at week 8 preferentially increases the low WT rE response such that both WT and SD*FL leading to comparable levels of NAb for both groups at week 12.

### Mapping DENV NAbs stimulated by WT and SD*FL rE vaccines

We used the week 12 high titer sera from the 3-dose study to further characterize NAbs stimulated by WT and SD*FL vaccine antigens. When the sera were tested for neutralization of the four DENV serotypes, we observed strong neutralization of DENV2 and no detectable neutralization of other serotypes demonstrating the dominance of DENV2 type-specific NAbs for both vaccine constructs (**Fig. 3A**).

**Figure 3.**
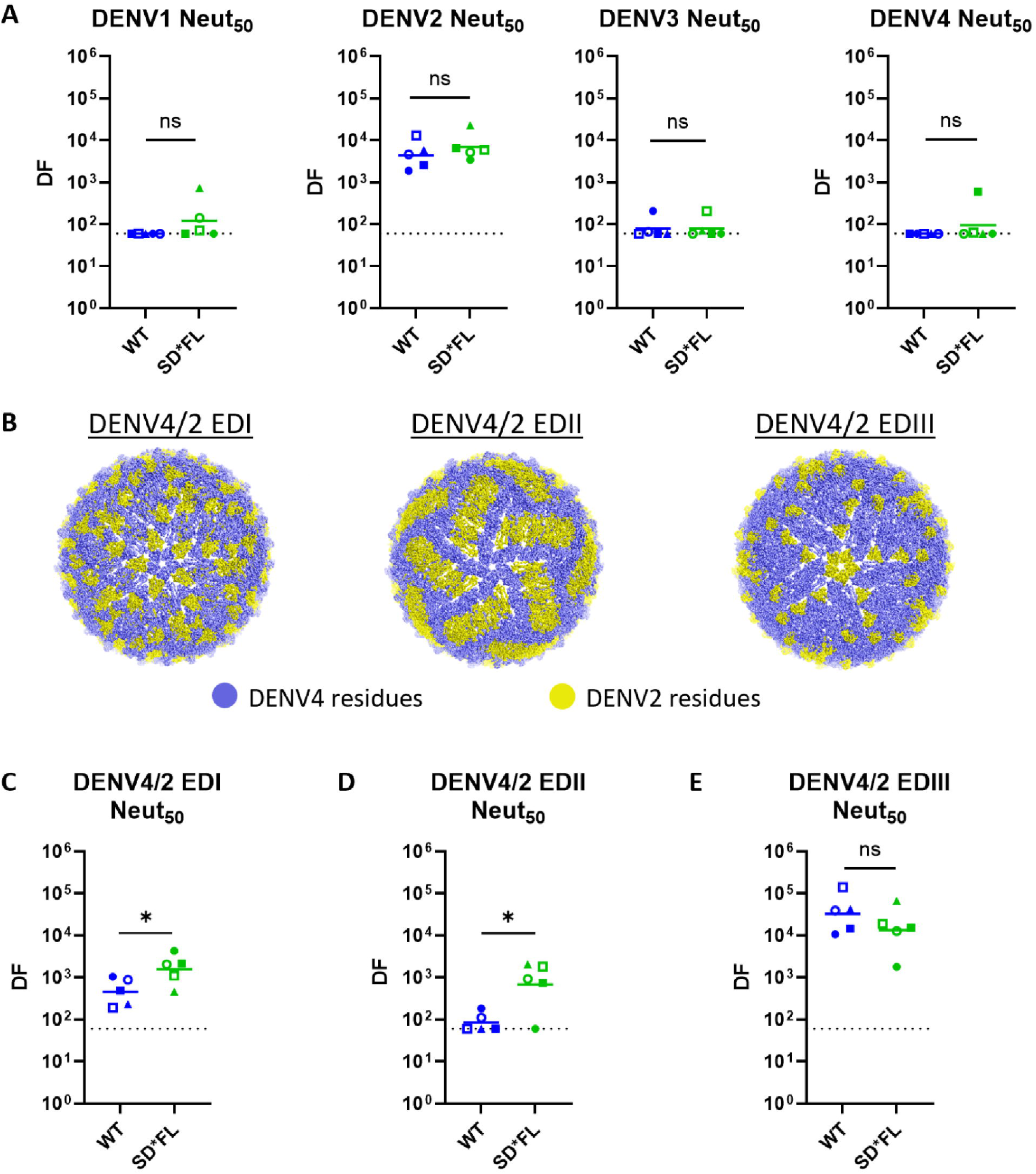
Mapping DENV2 E-domains targeted by neutralizing Ab using DENV4/2 chimeric viruses. (**A**) DENV1-4 50% neutralizing Ab titers at week 12. (**B**) DENV4/2 chimera schematic with DENV2 domain substitution highlighted in yellow (PBD 4CBF). The neutralizing IgG titer elicited by WT and SD*FL rE vaccines are reported as the Neut_50_ value against the parental viruses (**A**) DENV2, DENV4 and against the various domains of DENV2 on a DENV4 backbone (**C**) DENV4/2 EDI, (**D**) DENV4/2 EDII, (**E**) DENV4/2 EDIII. LoD for these assays is 1:60. The horizontal line represents the GMT of all mice within the group. Statistical analysis was done on log-transformed values, followed by unpaired t-test (two-tailed, p<0.05). The same mouse in different experiments is represented with the same color and symbols.

To further map vaccine sera, we used an existing panel of DENV4/2 E domain transplant viruses that have been previously used to map the epitopes of Abs from people exposed to WT DENV2 infections (**Fig. 3B**)(25). The chimeric viruses have DENV4 backbone with transplanted domain from DENV2 as indicated (EDI or EDII or EDIII) (**Fig. 3B**). Both WT and SD*FL antigens induced Abs that neutralize the DENV4/2 EDIII chimera demonstrating the presence of NAb targeting DENV2 EDIII, as well as Ab that spans EDII/EDIII such as 2D22(26) (**Fig. 3E**). SD*FL rE also elicited Abs that neutralized the DENV4/2 EDI and EDII chimeric viruses, whereas WT rE stimulated lower levels of NAbs that tracked with EDI (**Fig. 3C**) and no detectable NAbs to EDII (**Fig. 3D**). These results indicate that WT rE mainly induced EDIII targeting NAbs, while SD*FL rE stimulated a broader response directed to all three domains.

### Characterizing EDIII-Ab elicited by WT and stable dimer immunized mice

Antibodies that neutralize DENV2 have been mapped to simple epitopes contained within a domain or more complex epitopes that span two different domains on the same or different E molecules. For example, DENV2 type-specific and neutralizing mouse monoclonal Ab (mAb) 3H5 binds to a simple epitope on the lateral ridge of EDIII(27). In contrast, DENV2 type-specific and neutralizing human mAb 2D22 recognizes epitope centered on EDIII that expands into conserved amino acids on EDII of the second E protomer forming the homodimer. This cross-dimer interaction is critical for 2D22 binding to DENV2(20, 22). Consequently, unlike 3H5, 2D22 binds poorly to soluble EDIII (**Fig. S1**). We performed Ab depletion studies with recombinant DENV2 EDIII to measure the contribution of simple EDIII binding Abs to neutralization following vaccination with WT and SD*FL rE antigens (**Fig. 4A**). Sera were incubated with DENV2 EDIII coated beads and effective removal of EDIII binding antibodies was confirmed by ELISA (**Fig. 4B**). As expected, removal of simple EDIII binding Abs also led to a reduction in binding to E dimers (SD), which include simple EDIII epitopes (**Fig. 4C**). Next, we assessed how the removal of EDIII-binding Ab impacted DENV2 neutralization (**Fig. 4D**). In WT E immune sera, the removal of EDIII-binding Abs dropped DENV2 neutralization to a level below the detection limit of the assay. While the removal of EDIII-binding Abs also lowered NAb levels in SD*FL rE immunized animals, a sub-population of NAbs (∼30-40% of total response) was resistant to EDIII depletion (**Fig. 4E**). This result together with results obtained with the DENV4/2 chimeric E viruses suggests that SD*FL stimulates NAbs targeting all three domains on E protein.

**Figure 4.**
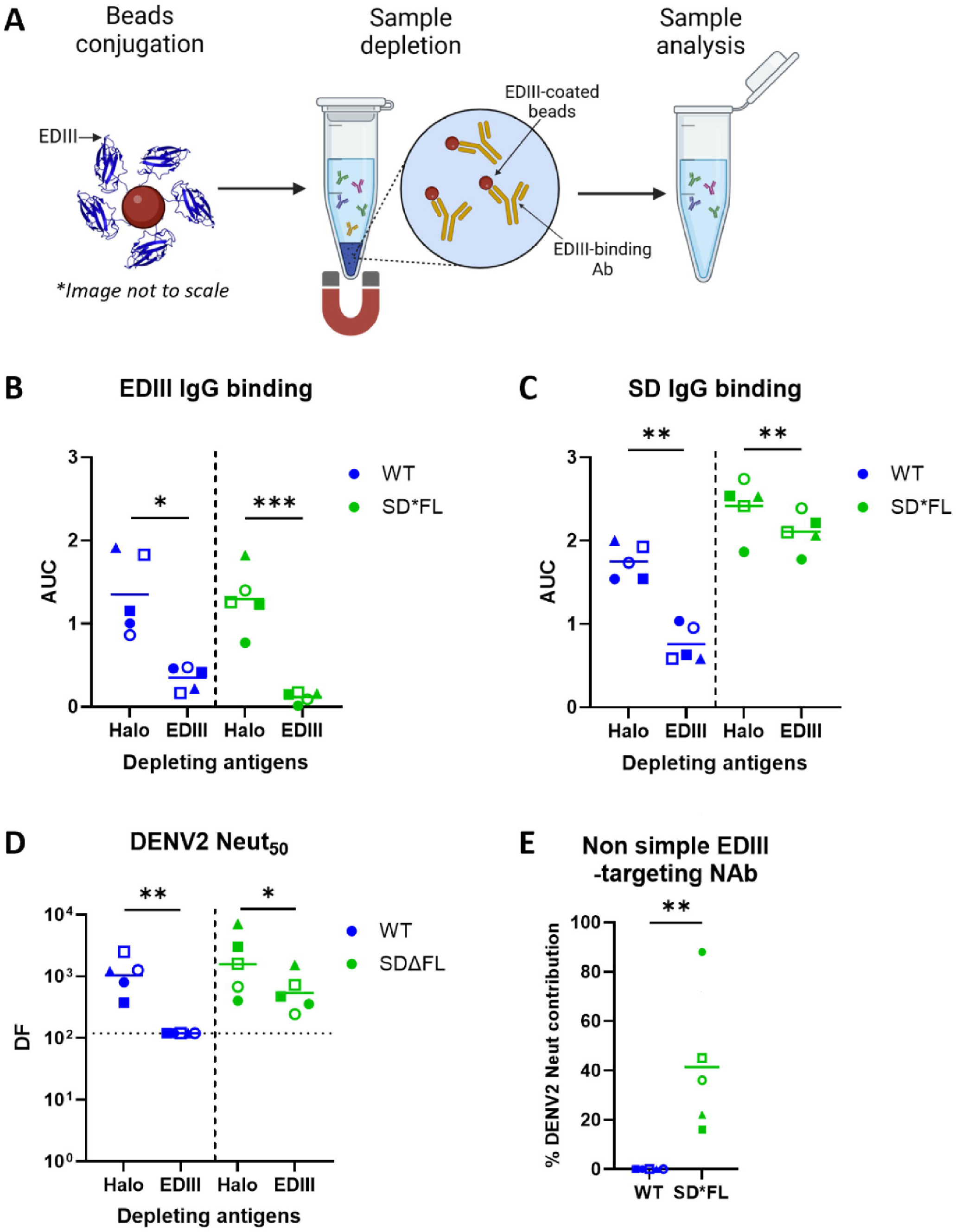
Characterization of EDIII binding antibody. (**A**) A schema for DENV2 EDIII depletion. First, we conjugated Magne HaloTag beads with either EDIII or HaloTag. Next, we treated the mouse sera with either EDIII- or HaloTag (control)-coated beads. Magnetic pulldown will separate any antibodies that bind to the coated beads, leaving behind unbound antibodies (non-simple EDIII or non-HaloTag antibodies) in the sera. Downstream depleted samples analysis include (**B**) DENV2 EDIII IgG binding, (**C**) DENV2 stabilized dimer with intact FL (SD) IgG binding, and (**D**) DENV2 neutralization, with LoD of DF120. (**E**) Percentage of NAb that target other areas outside of simple EDIII region. Each data point is the average of each sample’s duplicates. (**B, C, E**) The horizontal line represents the mean of all mice within the group. Statistical differences between Halo- vs EDIII-depleted samples (**B, C**) were determined by paired t-test (one-tailed, p<0.05). Unpaired t-test (one-tailed, p<0.05) was used to determine the statistical differences for (**E**). (**D**) The horizontal line represents the GMT of all mice within the group. Statistical analysis was done on log-transformed values, followed by paired t-test (one-tailed, p<0.05). The color and symbols match individual mice tested in different assays.

## Discussion

DENV protein subunit vaccines based on the ectodomain of E protein have not performed well in the past in non-human primates (NHPs) and human clinical studies(4, 5, 8, 9, 28). Vaccine stimulated NAb levels declined steeply over the first 6-12 months after vaccination (5–7). When vaccinated NHPs were challenged with DENV, some animals were protected, while others experienced breakthrough viremia(6, 7). While investigators had assumed that recombinant E protein produced as a soluble, secreted antigen formed homodimers similar to the protein on the viral surface, recent studies have demonstrated that the antigen is mostly present as a monomer at 37℃, which is a plausible explanation for the overall poor performance of DENV subunit vaccines(8, 9). The main goal of our study was to compare the properties of DENV2 NAbs induced by soluble E protein antigens that are monomers (WT) or homodimers (SD*FL) at 37℃. Our results demonstrate that stable homodimers stimulated DENV2 NAbs targeting multiple regions of the viral envelope, whereas the WT E antigen stimulated NAbs that mainly target simple EDIII epitopes. Together, our results demonstrate that by altering the oligomeric state of DENV2 E protein to better match the antigen on the surface of the infectious virus, it is possible to elicit Abs against all three domains of the E protein.

Currently, the leading DENV vaccine candidates are based on tetravalent live attenuated flavivirus platforms. These vaccines have been challenging to develop because of inherent replication differences between vaccine viruses resulting in unbalanced immunity skewed to a single serotype and/or a single genotype within a serotype(15, 29, 30). In clinical trials, immunodominance of a single vaccine serotype over the other three serotypes has been linked to poor outcomes, including vaccine-enhanced WT DENV infections(16). Given these challenges with LAVs, a protein subunit vaccine that mimics the architecture of the viral envelope is a promising alternative strategy that is not dependent on balanced replication of each vaccine component.

Investigators have relied on DENV neutralization assays both to characterize vaccine responses and as a correlate of protection. However, in clinical trials, the presence of NAbs to each serotype alone was not a reliable predictor of protection. Recent studies have indicate that DENVs produced for laboratory studies are inefficiently processed and mostly consist of partially mature virions(31). These partially mature virions are hypersensitive to neutralization by some low-affinity antibodies, which are unlikely to be protective against mature DENVs circulating in humans(31). To control for virus maturation state and selectively measure vaccine responses that are effective against fully mature DENV2 virions, we used fully mature DENV2 stocks produced in a cell line overexpressing a protease necessary for DENV processing and maturation(32). Under these conditions, we observed that 2 doses of the stabilized E homodimer stimulated higher levels of mature DENV2 NAb compared to WT E protein. These results are consistent with our previous work(10) demonstrating that 2 doses of the stabilized E homodimer stimulates higher levels of NAb than WT rE in mice, although the original study did not control for DENV maturation state. In the current study, DENV2 NAb titers were comparable in the 3-dose study indicating that the low response stimulated by 2 doses of WT rE is preferentially boosted by the third dose. The animals were only monitored for 4 weeks after the third dose and we have not addressed durability of NAbs in this study.

Neutralizing Abs that target different regions of the E-protein would be able to block viral entry at different stages (attachment, internalization, membrane fusion) and minimize vaccine escape. The different domains of the DENV E-protein play different roles in various stages of the virus lifecycle. EDIII is involved in the initial virus attachment with host cell receptors(33). EDII is critical for the dimerization of E protein, and contains the fusion loop required for viral and host cell membrane fusion during viral entry into cells(33, 34). People exposed to DENV infections develop NAbs to multiple epitopes on different E protein domains, and these responses that are broadly effective against variant genotypes belong to a serotype(22, 25, 39, 40). In contrast, live attenuated DENV vaccines appear to stimulate a more focused NAb response with a narrow breadth across genotypes within a serotype, with the potential for breakthrough infections with vaccine mispatched genotypes(29, 30, 39, 41). In the current study, the WT E vaccine stimulated a focused NAb response that was mainly directed to EDIII. Previous studies have demonstrated that the lateral ridge region of EDIII contains epitopes targeted by some type-specific NAbs (35, 36). However, these epitopes are variable between DENV genotypes belonging to the same serotype and responsible for variable neutralization potency including escape from EDIII antibodies(35, 37, 38). Our studies showed that the SD*FL vaccine elicited a NAb response that targeted epitopes on all three domains. While a NAb response to multiple targets on E protein is likely to have more breadth than a response focused to a single site, follow up studies are needed to directly assess efficacy of WT and SD*FL rE vaccines against vaccine matched and variant DENV2 genotypes.

In conclusion, our studies support further development and evaluation of E dimer-based subunit vaccines, demonstrating that DENV2 stable homodimers stimulate high levels of NAbs targeting multiple domains on E protein. We have found that the mutations used for stabilizing DENV2 homodimers also work for the other three serotypes and Zika virus with minor virus-specific modifications(11). Subunit vaccine studies are in progress with DENV1, DENV3 and DENV4 because DENV vaccines must be tetravalent to minimize the risk of ADE. Moreover, we are evaluating different vaccine delivery platforms (nanoparticles, liposomes, nucleic acid) and adjuvants to increase the immunogenicity and durability of vaccine responses.

## Methods

### Cell lines and viruses

EXPI293F (Thermo Fisher Scientific) cells were maintained in EXPI293 expression medium at 37°C with 8% CO_2_ at 250 rpm.

Vero-81 (ATCC CCL-81) cells were maintained at 37°C with 5% CO_2_ in DMEM-F12 (Gibco) supplemented with 5% of heat-inactivated fetal bovine serum (HI FBS, Gibco), 1% of Penicillin-Streptomycin (Gibco), 1% L-Glutamine (Gibco), and 1% MEM non-essential amino acids (MEM NEAA, Gibco).

<u>Vero-furin (VF) cells</u> were given by the Baric lab at UNC Chapel Hill(32). The cells were maintained at 37°C with 5% CO_2_ in DMEM-F12 + 5% HI FBS + 1% Anti-anti (Gibco) + 1% MEM NEAA + 1% L-Glutamine.

#### Viruses

All DENV2 S16681 used in the experiments were grown in VF cells, except for DENV2 S16681 used in the 2-dose study ELISA which was grown in Vero cells. For 3-dose study ELISA, DENV2 S16681 VF-grown virus was concentrated using Centricon® Plus-70 Centrifugal Filter (Millipore) per manufacturer instruction to obtain the amount and concentration of virus required for ELISA.

Cross-neutralization assessment was done using DENV1 WestPac-74, DENV3 CH53489, DENV4 Sri Lanka 92. All viruses were grown in VF cells.

Chimera viruses and EDIII depletion experiments used the following viruses: DENV2 16803, DENV4 Sri Lanka 92, DENV4/2 EDIII, DENV4/2 EDII and DENV4/2 EDI that were grown using a previously described method(25). All viruses were grown in VF cells. As the chimeric panel is based on DENV2 strain S16803, we used DENV2 S16803 for our mapping study. DENV2 strains 16681 and 16803 E protein sequences differ at 5 positions (M6I, R120T, I141V, I164V, and E383D) – all are outside of the SD*FL mutations.

### DENV2 sE plasmid generation

Cloning method for various DENV2 S16681 variants sE into PαH mammalian expression vector was described previously(10). For the HisAvi-variants, after the C-terminal His8-tag, glycine linkers (GGCAGCTCTGGCGGCAGC) and AviTag (GGCCTGAACGATATTTTCGAGGCCCAGAAAATCGAGTGGCACGAGTAG) were added. The DENV2 sE cloned plasmids were heat-shock transformed into *Escherichia coli* DH5α cells for amplification, miniprepped or midiprepped using DNA endotoxin-free kits, and stored in endotoxin-free H_2_O at −20°C (Macherey-Nagel).

### DENV2 sE protein large-scale expression

#### For DENV2 WT-His

Stable line was grown up in Expi293 supplemented with 250 µg/ml geneticin, with geneticin removed during the final amplification step. The cultures were supplemented with 5 ml of 100X GlutaMAX (Thermo 35050061) and 50 ml of 6% HyClone Cell Boost Boost I (Cytiva SH30584.02) in OptiMEM (Thermo 31985070) per liter of culture when the cells reached a density of approximately 1.5×10^6^ cells/ml (day 4 after reaching the final culture scale). The cultures were harvested when the cells dropped below 90% viability on day 9 after reaching the final scale, with the centrifugation, filtering, and purifications done the same as for the proteins made in Expi293 transfection cultures.

#### For other DENV2 variants-His

Expi293 cells were transfected per manufacturer instructions (Thermo Scientific), using 1mg of DNA/1L of culture volume for each flask. Enhancers 1 and 2 were added on day 1 post-transfection, then the cultures were harvested on day 3 at 12,000 x g for 1 hour at 4°C. The supernatants were filtered (0.22 µm), then stored at 4°C until purification.

#### For biotinylated SD-HisAvi

Expi293 cells were co-transfected per manufacturer instructions (Thermo Scientific) in 1L volume with 900 µg of a plasmid with SD-HisAvi (UNC Protein Core) and 100 µg of mammalian BirA ligase expression plasmids (UNC Protein Core). Culture media was supplemented with 100 µM biotin before transfection to aid SD-HisAvi biotinylation by BirA ligase. Downstream processes (enhancers and harvest timeline) are the same as DENV2 variants-His.

### DENV2 sE protein purification and analysis

Filtered supernatants were buffer-exchanged via tangential-flow filtration (10 kDa MWCO) into cobalt-binding buffer (50 mM sodium phosphate pH 7.4, 500 mM NaCl, 5 mM imidazole, 0.02% sodium azide, 10% glycerol), with 4 total buffer exchanges. The supernatant was then loaded at 1.5 ml/min recirculating overnight with a peristaltic pump and flow adapter onto a hand-packed TALON column. The next day, the column was switched to flow-through, and once completely loaded, the column was washed with 10 Column Volume (CV) binding buffer. After a 10 CV second wash with binding buffer supplemented with 10 mM imidazole, bound proteins were eluted in two steps (one of 5 CV, and one of 2.5 CV) with binding buffer supplemented with 150 mM imidazole. Column fractions were assessed by SDS-PAGE and fractions containing target protein were pooled, filtered, and concentrated before loading onto a Superdex 200 16/600 column equilibrated in 1x PBS + 10% glycerol. The sizing column fractions were assessed by SDS-PAGE, and DENV2 protein-containing fractions were pooled, filtered, and concentrated before aliquoting and flash-freezing in liquid nitrogen. The endotoxin level in the final stock was measured using Endosafe-PTS cartridges (#PTS20F; Charles River) on a nexgen-PTS reader between 1:20 - 1:40 dilution in endotoxin-free water.

#### For biotinylated SD-HisAvi

To assess the success of the biotinylation reaction, column fractions were assessed by SDS-PAGE, with elution fractions assessed for biotinylated protein by mixing with a 2x molar excess of avidin. Downstream processes are the same as above.

### DENV2 EDIII- & HaloTag-His production

A Halo-tag fused DENV-2 EDIII (ADA00411·1, E protein aa 297-394) was expressed in Expi293 with an N-terminal human serum albumin secretion signal peptide and a C-terminal His6-tag as previously described(42). A Halo-tag construct, lacking DENV-2 EDIII, was also produced similarly with a C-terminal His6-tag. The DENV2 EDIII and Halo-tag proteins were purified using Ni-NTA agarose (Qiagen) from the cell culture medium. Purified DENV2 EDIII was biotinylated by incubating 100 µg of DENV2 EDIII with 5 nmol of the HaloTag PEG biotin ligand (Promega) at room temperature (RT) for 2 hours. Excess biotin ligand was removed by desalting column. Biotinylated DENV2 EDIII was assessed by comparing the mobility of the biotinylated EDIII in the presence and absence of streptavidin in SDS-PAGE.

### Mouse immunizations

Female Balb/c mice were purchased from Jackson Laboratory and used in immunization studies at 6 weeks of age. Mice were injected subcutaneously in the flank with 5 µg DENV2 rE subunit vaccine antigen in 100 µL PBS. Vaccine groups (n = 5) included placebo (PBS), WT rE, WT rE + Alum, SD*FL rE, SD*FL rE + Alum. Adjuvant Alum (Alhydrogel, InvivoGen) was included at 125 μg per dose. Mice were immunized twice with the same antigen formulation and dose level at days 0 and 21. In another experiment, a third dose was given at day 56. Serum samples were collected at various time points as indicated for antibody analysis.

### Ethics statement

All experiments involving mice were performed according to the animal use protocol approved by the University of North Carolina Animal Care and Use Committee. The animal care and use related to this work complied with federal regulations: the Public Health Service Policy on Humane Care and Use of Laboratory Animals, Animal Welfare Act, and followed the Guide for the Care and Use of Laboratory Animals.

### Enzyme-linked immunosorbent assay (ELISA)

#### Whole virus ELISA

DENV2 IgG binding titer was measured by DENV2 antigen capture ELISA. A high-binding 96-well plate (Greiner bio-one) was coated using 0.2 μg/ml of 1M7 (DENV fusion loop human mAb) in 0.1M bicarbonate buffer (pH 9.6) overnight at 4°C, and washed with wash buffer (1X Tris-buffered saline (TBS) + 0.2% Tween-20). The plate was blocked with blocking buffer (3% non-fat dried milk in 1X TBS + 0.05% Tween20), before DENV2 was loaded to each well. Next, serially diluted mouse sera were added onto the plate. Plates were washed in between each step, and each incubation was done for 1 hour at 37°C. Bound antibodies were detected by alkaline phosphatase (AP)-conjugated anti-mouse IgG (1:1000, Sigma-Aldrich) for 45 min at 37°C, and washed. Plates were developed using AP substrate (SIGMAFAST p-Nitrophenyl phosphate tablet, Sigma-Aldrich) and measured at 405 nm. IgG titer was determined by area under the curve (AUC) in GraphPad Prism using nonlinear fit – sigmoidal dose-response (variable slope).

#### Soluble E-coated ELISA

Nickel-coated ELISA plates (Thermo Fisher Scientific, Pierce 15142) were used to capture 0.2 μg/ml of rE protein in 1X TBS for 1 hour at 37°C. The plate was washed with wash buffer, and serially diluted sera in blocking buffer were added for 1 hour at 37°C and subsequently washed with wash buffer. Downstream processes are the same as above for whole virus ELISA.

#### Biotinylated-protein ELISA

High-binding 96-well plate was coated using 0.2 μg/ml of Streptavidin (Invitrogen) in 1X TBS overnight at 4°C or 1 hour at 37°C. Supernatant was removed, and 0.2 μg/ml of biotinylated antigens (DENV2 EDIII or SD) was added in 1X TBS, and incubated for 1 hour at 37°C. The plate was washed using wash buffer and then blocked with blocking buffer for 1 hour at 37°C. The plate was washed, and serially diluted sera in 1X TBS were added for 1 hour at 37°C and subsequently washed with wash buffer. Bound antibodies were detected by AP-conjugated anti-mouse IgG (1:1000 in 1X TBS) for 45 min at 37°C, and washed. Plates were developed using AP substrate and measured at 405 nm. IgG titer was determined by area under the curve (AUC) in GraphPad Prism using nonlinear fit – Sigmoidal, 4PL.

### Focus reduction neutralization (FRNT) assay

#### Cell seeding

Vero cells were seeded in 96-well plate at 2×10^5^cells/ml in 100ul of cell culture media per well. Plates were incubated overnight at 37°C.

#### Cell infection

Sera were serially diluted in infection media (similar to cell culture media, only with 2% HI FBS) and incubated with a fixed amount of virus at 1:1 v/v ratio for 1 hour at 37°C. The amount of virus had been previously determined to give a minimum of 30 foci/well. In each experiment, the starting serum dilution was different and based on the NAb titer of the specimen and the quantity of sample remaining. The limit of detection (starting dilution) for Fig. 1G and 2F is 1:10, Fig. 3 is 1:60, Fig. 4D is 1:120. Culture media on cells were removed, and serum/virus complex was added onto cells for 1 hour at 37°C. Serum/virus complex was removed from cells, and overlay media (OptiMEM (Gibco) + 2% HI FBS + 1% anti-anti + 1% (w/v) carboxymethylcellulose (Sigma-Aldrich)) was added to cells. Cells were left to incubate for 42-48 hours (depending on the viruses used).

#### Cell harvest and staining

Cells were washed with 1X phosphate-buffered saline (PBS), and fixed with 4% paraformaldehyde. Next, cells were permeabilized with 1X permeabilization (Perm) buffer (10X Perm buffer was diluted in MilliQ water. 10X Perm: 1% of bovine serum albumin (BSA, Sigma), 1% of Saponin (Sigma), 0.1% of sodium azide (Sigma) in 10X PBS), and blocked using blocking buffer (5% non-fat dry milk in 1X Perm) at RT for 10 min. Blocking buffer was removed, and cells were stained using 1M7 at 1.2×10^-3^mg/ml for 1 hour at 37°C. Cells were washed with 1X PBS, and stained with mouse anti-human IgG Fc HRP (Southern biotech) at 1:4000 for 1 hour at 37°C. Cells were washed and stained with KPL TrueBlue Peroxidase Substrate (SeraCare). Cells were then rinsed with MilliQ water and plates were left to dry prior to imaging with ImmunoSpot S6 analyzer (Cellular Technology Limited). Foci were counted using Viridot on RStudio, and checked for accuracy(43). Neutralization curves were generated by plotting sera dilution factor (log) vs % neutralization. Curves were fitted using nonlinear fit – sigmoidal dose-response (variable slope) with constraints (top = 100, bottom = 0). Curves were acceptable when R^2^ ≥0.75 and Hill Slope >0.5. Neut_50_ value is the dilution factor at which sera neutralize 50% of the virus. Analysis was done using Graphpad Prism

##### DENVs chimera neutralization

All steps are identical to FRNT assay explained above, with the exception of serum/virus complex was not removed prior to the addition of overlay media.

### EDIII Depletion

#### Protein-beads conjugation

First, Magne HaloTag beads (Promega) were transferred into Lo-bind tubes (Eppendorf), and beads were washed with 1X TBS for a total of 3 times. To separate beads from supernatant, magnetic stand (DynaMag2, Invitrogen) was used. After the third wash, pre-determined amount of biotinylated protein was added to the beads. Ratio of initial beads volume (μl) to antigen amount (μg) is 1:2. Beads and biotinylated protein were left to incubate for 2 hour at RT, mixing at 1300rpm in ThermoMixer C. Once the conjugation was done, beads were washed 3 times using blocking buffer (5% BSA in 1X TBS). Beads were further blocked using blocking buffer for 1 hour at RT, mixing at 1300rpm. After blocking step, beads were washed 3 times with 1X TBS and re-suspend in 1X TBS. The final resuspension volume is equal to the total volume of all diluted sera that is going to be depleted; the final protein concentration in this resuspension is 48 μg/ml. Beads were stored at 4°C in lo-bind tubes if not being used immediately.

#### Sera depletion

Depletion was done using non-binding 96-well plate (Corning). Volume of conjugated beads/well is equal to volume of depleted sera/well. Conjugated beads were added to each well, and the supernatant was discarded by placing the plate on 96-well plate magnetic separator (Millipore Sigma). Next, diluted sera (1:60, in 1X TBS) was added per well and plate was covered with adhesive film (Thermo). The plate was incubated at 37°C for 1 hour at 800rpm. Depletion was done for 2-3 rounds. Sera depletion was considered complete when EDIII-depleted sera (1:1200) against EDIII antigen on an ELISA reached background level compared to undepleted sera. Depleted sera were stored in untreated polypropylene 96-well plates (Corning) at 4°C.

BioRender was used to make schemas for mouse vaccination study and EDIII depletion.

## Data availability

not applicable

## Acknowledgment

We thank Edwing D. C. Cuadra for discussions and suggestions with the EDIII depletion experiments.

