## Supplemental Table and Figure for "Stabilized dengue virus 2 envelope subunit vaccine redirects the neutralizing antibody response to all E-domains"

**Supplementary Table 1.** DENV2 Soluble E proteins (Name and mutations)used as vaccines and/or in immune assays. Structures based on PDB 1OAN.

| | WT | Monomer/ M2P4 | SD | SD*FL |
|---|---|---|---|---|
| Mutations | | M2 (G258E), P4 (S29K, T33V, A35M) | I2 (A259W, T262R), U6 (F279W, T280P) | I2, U6, I8 (G106D) |
| Previously published name[10] | | SC.25 | SC.14 | SC.10 |
| | 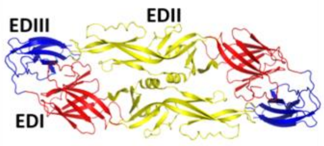  | 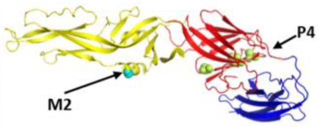  | 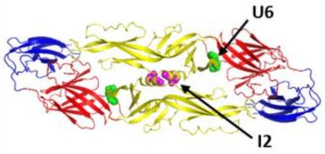  | 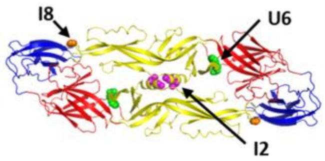  |

### Supplementary Figure 1

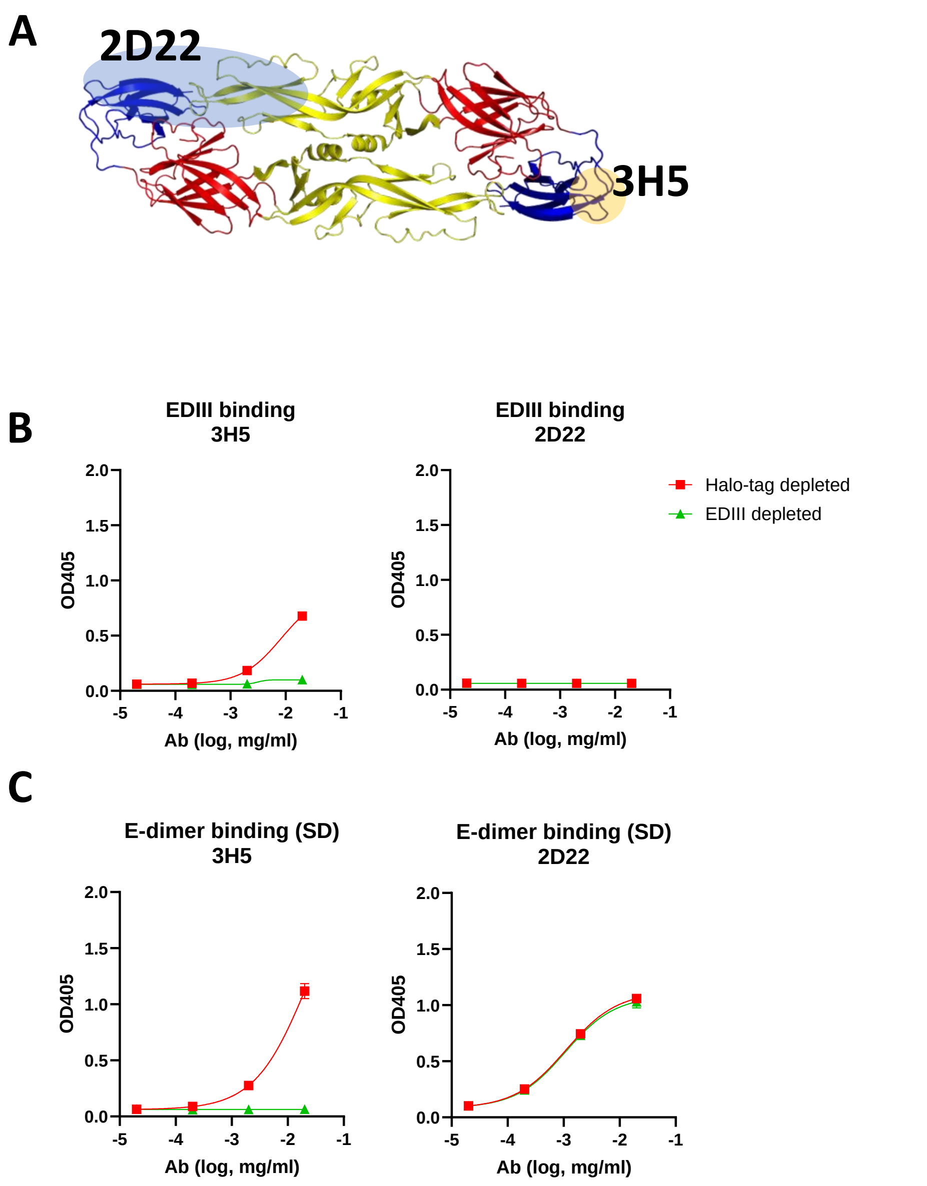


**Supplementary Figure 1. Recombinant EDIII antigen depletes Abs binding to simple EDIII epitopes but not Abs binding to quaternary structure epitopes that include EDIII** (**A**) Epitope location of simple EDIII-targeting MAb 3H5 and quaternary epitope-targeting MAb 2D22. The MAbs were incubated with immobilized recombinant EDIII or a control antigen (Halo-tag) to deplete EDIII binding Ab and tested for binding to (**B**) EDIII or (**C**) E-dimer (SD).
